# Evolution of ovipositor length in *Drosophila suzukii* is driven by enhanced cell size expansion and anisotropic tissue reorganization

**DOI:** 10.1101/466375

**Authors:** Jack E. Green, Matthieu Cavey, Emmanuelle Caturegli, Nicolas Gompel, Benjamin Prud’homme

## Abstract

Morphological diversity is dominated by variation in body proportion. Yet the cellular processes underlying differential growth of morphological traits between species remain largely unknown. Here we compare the ovipositors of two closely related species, *Drosophila melanogaster* and *D. suzukii*. *D. suzukii* has switched its egg-laying niche from rotting to ripe fruit. Along with this shift, the *D. suzukii* ovipositor has undergone a significant change in size and shape. Using an allometric approach we find that, while adult ovipositor width has hardly changed between the species, *D. suzukii* ovipositor length is almost double that of *D. melanogaster*. We show that this size difference mostly arises during a 6-hour time window in the middle of pupal development. We observe that the developing ovipositors of the two species comprise an almost identical number of cells, with a very similar profile of cell shapes and orientations. After cell division stops, we find that the ovipositor area continues to grow through the isotropic expansion of cell apical area. Remarkably, at one point, the rate of cell apical area expansion is more than 4 times faster in *D. suzukii* than in *D. melanogaster*. In addition, we find that an anisotropic cellular reorganization of the developing ovipositor results in a net elongation of the tissue, despite the isotropic expansion of cell size, and is enhanced in *D. suzukii*. Therefore, the quantitative fine-tuning of shared, morphogenetic processes -the rate of cell size expansion and the cellular rearrangements–can drive macroscopic evolutionary changes in organ size and shape.

## Introduction

Animal morphological diversity results from changes in the position, number, colour, size and shape of body parts. Developmental and evolutionary geneticists have extensively studied the changes in the position, number or colour of body parts (1–5). In contrast, the mechanisms underpinning the evolutionary changes in body part size and shape have received much less attention, despite the fact that these constitute an enormous source of diversity (6). The pioneering work of D’Arcy Thompson (7) and Julian Huxley (8) described changes in body part size and shape with scaling relationships and mathematical equations. How these scaling relationships between species translate at the cellular and tissue levels remains, however, poorly characterized (9–12). A major challenge now is to understand how changes in size and shape of morphological traits between species can be explained in terms of cell proliferation, size and shape, cellular interactions and organisation, and the physical forces acting within and between cells and tissues (13–15).

To bridge the gap between the mechanisms sculpting morphogenesis during development and the mechanisms driving variation in body part size and shape during evolution, we investigate two closely related *Drosophila* species, *D. melanogaster* and *D. suzukii*. *D. suzukii* is an invasive pest species that lays its eggs in a range of ripe fruits, causing substantial economic losses (16). By contrast, *D. melanogaster*, like most Drosophila species, lays eggs almost exclusively in rotting fruit (17). This ecological shift between these species is accompanied by changes in size and shape of the egg-laying organ, or ovipositor, which is used to pierce and dig into selected egg-laying substrates. *D. suzukii* has evolved an enlarged and serrated ovipositor that enables the females to penetrate and lay eggs inside the tougher skin of ripe fruit (18). Our aim here is to compare the development of the *D. suzukii* and *D. melanogaster* ovipositors in order to identify the cellular processes responsible for the quantitative changes in size and shape observed in the adult structures of these species.

## Results

### The *D. suzukii* ovipositor is almost twice as long as that of *D. melanogaster*

Along with a novel egg-laying preference for ripe fruit, *D. suzukii* has evolved an enlarged ovipositor compared with its close relatives (Fig. 1A). The dimensions of the ovipositor can be measured to a first approximation as a flattened plate (Fig. 1B, C). There is a marginally significant, small difference in ovipositor width between the species (*D. melanogaster* width = 48±1 μm; *D. suzukii* width = 52±1 μm; Student’s t-test p = 0.02). Nevertheless, the major size difference is driven by the ~60% increase in ovipositor length in *D. suzukii* (*D. melanogaster* length = 261±2 μm; *D. suzukii* length = 414±4 μm; Student’s t-test p < 0.001). (Raw data is provided in Supplementary Table 1). However, the length of the ovipositor most likely scales with overall body size and since body size can vary between individuals and between species, we suspected that such variation might obscure a simple interspecific comparison of ovipositor length. We therefore used an allometric approach to characterize the difference in the length of the adult ovipositor between *D. melanogaster* and *D. suzukii* in greater detail.

**Fig 1.**
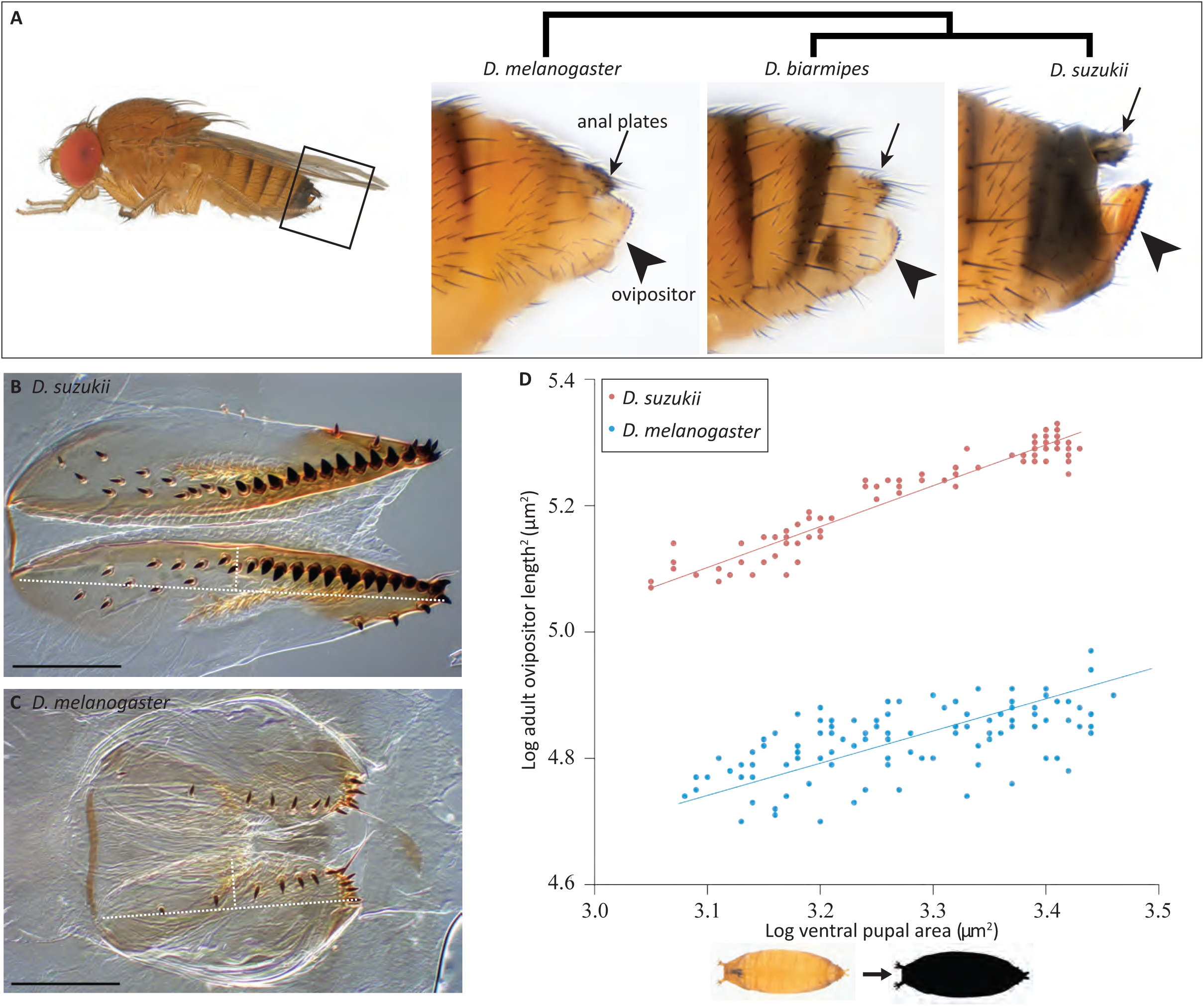
Evolutionary shift in the scaling relationship of ovipositor length against body size between *D. suzukii* and *D. melanogaster*. (A) Adult ovipositors of 3 closely related *Drosophila* species, *D. melanogaster*, *D. biarmipes* and *D. suzukii*, in lateral profile (arrowhead indicates ovipositor, arrow indicates anal plates). *D. suzukii* female on the left; boxed area indicates the approximate posterior region shown in the panels. Images reproduced from (18). (B, C) *D. suzukii* and *D. melanogaster* adult ovipositors, respectively. The long, white, dashed line indicates the measured length; the short indicates half width. Scale bar is 200 μm. (D) Scaling relationship of ovipositor length squared and body size in *D. melanogaster* (blue; n=114) and *D. suzukii* (red; n=99) on a log-log scale. Overall body size is measured using ventral pupal area as a proxy, as illustrated on the x-axis. The slope is modestly steeper in *D. suzukii*, but more importantly, the intercept is shifted upwards, indicating that ovipositor length is enlarged across the full range of body sizes.

Allometry refers to the phenomenon of biological scaling, namely the universal property of morphological traits to scale with overall body size (8, 19). For the adults of both species, we plotted nutritional allometries – the diet of the flies was systematically manipulated to generate the full range of viable adult body sizes. This gave us a complete description of how ovipositor length covaries with body size, and so enables a more robust comparison of the scaling relationship between *D. melanogaster* and *D. suzukii*.

For both species, the scaling relationship of ovipositor length squared and ventral pupal area is linear when plotted on a log-log scale, and so can be described by a linear function with a slope and intercept (20, 21) (Fig. 1D). The slope of the scaling relationship is steeper in *D.* suzukii, increasing by 27% (*D. melanogaster* slope = 0.51, 95% C.I. = 0.44-0.59; *D. suzukii* slope = 0.65, 95% C.I. = 0.61-69; common slope test p < 0.01). This implies that the mechanism that coordinates changes in nutrition with ovipositor growth has diverged between species. Nevertheless, in both species, the ovipositor is a hypoallometric trait (slope < 1) – that is to say, within a species, ovipositor size is relatively invariant across the body size range.

Furthermore, it is clear that the *D. suzukii* ovipositor is proportionally nearly twice as long as the *D. melanogaster* ovipositor, across all body sizes (Fig. 1D). Therefore the mechanism that determines the final ovipositor length for any given body size has diverged between species. We next sought to establish the growth trajectory of the ovipositor through developmental time in both species. In particular, by comparing the trajectories, we wanted to determine when the size difference in the ovipositor appears, and whether it appears gradually or suddenly in time.

### The ovipositor primordium does not differ in size by the end of larval development

The vast bulk of imaginal disc growth occurs during the third larval instar and we therefore focussed our efforts on this stage. We measured overall genital disc area, and the apical area of ventral cells that include the primordium of the external genitalia (22). For this, we used an antibody against E-cadherin to label adherens junctions, and therefore apical cell membranes. Using the overall disc area and ventral cell size measurements, we estimated ventral cell number. Larval development is prolonged in *D. suzukii* by approximately one day, and so while the absolute rate of disc growth is slower in *D. suzukii*, the duration of the growth period is extended in compensation (Fig. S1A, B). After taking this into account, the relative growth trajectories of the female genital disc are essentially indistinguishable during the third larval instar between *D. melanogaster* and *D. suzukii* for both overall genital disc area and ventral cell number (Fig. S1C, D). Although the cell size trajectories differ somewhat between the species, ultimately, we find no significant difference in the apical area of ventral cells by the wandering stage (Fig. S1E; *D. melanogaster* = 7.13±0.24 μm; *D. suzukii* = 7.87±0.64 μm; Student’s t-test p > 0.05).

Although the genital discs of *D. suzukii* and *D. melanogaster* contain the same number of cells, the relative size of the ovipositor primordium within the genital disc might have changed between species. Previous fate maps of the female genital disc have established that anterior-ventral cells give rise to the internal genitalia and are marked by *abdominal-A* (*abd*-*A*) expression (Fig. S1F) (22, 23) and posterior-ventral cells give rise to the external genitalia, but express no known marker. We discovered that the expression of the gene *teashirt* (*tsh*) is restricted to a population of posterior-ventral cells (Fig. S1F, G, J). Moreover, it has a sharp boundary and a mutually exclusive expression domain with *abd*-*A* (Fig. S1H, K). In addition, we find that *tsh* expression overlaps with the expression of a Gal4 line (19D09-Gal4) that marks the ovipositor fate from larval stages to adult in *D. melanogaster* (Fig. S1F, I, M-O). The mutual exclusion of the *tsh* and *abd*-*A* expression patterns and the agreement between the *tsh* and 19D09-Gal4 expression support our interpretation that *tsh* labels the future external genitalia (Fig. S1F).

Examining the overlap between 19D09-Gal4 and *tsh* expression in *D. melanogaster*, we find that the ovipositor primordium lies entirely within the *tsh*-positive domain, and indeed represents the majority (~70%) of the *tsh*-positive domain by area (Fig. S1I). Hence we can use the *tsh* expression domain as a reasonable proxy for the ovipositor primordium. We find no significant difference in the relative area of the *tsh*-positive territory between the species at wandering stage (Fig. S1L). This strongly suggests that the ovipositor primordia are the same size in *D. suzukii* and *D. melanogaster* by the end of larval development.

In summary, the interspecific difference in ovipositor size has not appeared by the end of larval development. We therefore turned our attention to ovipositor development during metamorphosis.&

### The interspecific difference in ovipositor size and shape is generated between 48 and 54 hours in pupal development

To address when the interspecific size difference first appears, we needed a set of markers to demarcate the ovipositor in a comparable way, such that its development could be tracked in both species. Unfortunately, *tsh* expression in the presumptive external genitalia disappears by ~12 h APF (hours after puparium formation). However, between 18 and ~30 h APF, we found that the gene *senseless* (*sens*) is expressed in a row of discrete cells in the presumptive ovipositor in both species, most likely the bristles precursor cells (24) (Fig. 2A-F). The transitions in *senseless* expression and the occurrence of key morphogenetic events (described below) take place at approximately the same absolute time points after puparium formation in both species. Prominent amongst these key morphogenetic events is a sequence of changes in the shape of the egg-laying cavity from 18 to 30 h APF. It changes from a narrow triangular slit (~18 h APF; Fig. 2A, D), to a broader keyhole-like hollow (~24 h APF; Fig. 2B, E) and finally to a thinner and more elongated cavity (~30 h APF; Fig. 2C, F). Soon after 30 h APF, the presumptive ovipositor starts to project out and adopt the form of a blade (Fig. 2G, J). The blade continues to elongate from 36 to 54 h APF in both species (Fig. 2G-L), and can be distinguished from surrounding tissue on morphological criteria alone.

**Fig. 2.**
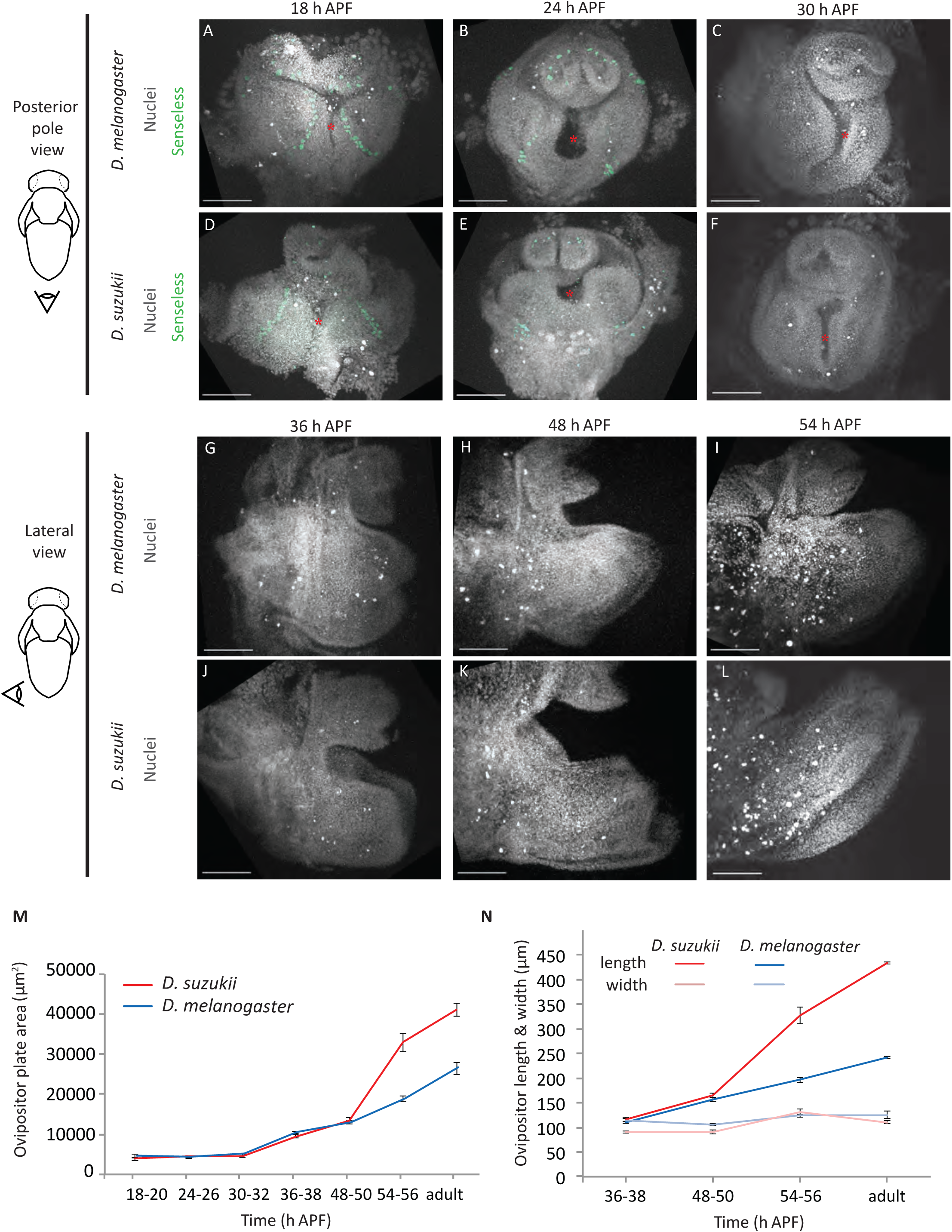
The evolutionary divergence in ovipositor size and shape is generated in a restricted time window during pupal development. (A-F) Ovipositor development from 18 to 30 h APF in (A-C) *D. melanogaster* and (D-F) *D. suzukii*, respectively. The presumptive ovipositor plates are arranged as a pair of lobes on either side of the future egg-laying cavity. (G-L) Ovipositor development from 36 to 54 h APF in (G-I) *D. melanogaster* and (J-L) *D. suzukii*, respectively. The presumptive ovipositor projects out and elongates over this period. (A-L) All images are maximum projections of confocal stacks, with nuclei shown in grey and Senseless expression in green in (A-F). A red asterisk indicates the egg-laying cavity. Schematic on the left illustrates the image orientation with respect to the pupa; images are posterior (A-F) or lateral views (G-L). All images are to the same scale; scale bar is 50 μm. (M) Growth in the mean, total ovipositor plate area during metamorphosis, in *D. melanogaster* (blue) and *D. suzukii* (red). Number of samples at the 7 time points in chronological order, in *D. melanogaster*: n=8, 13, 9, 10, 10, 10, 10; in *D. suzukii*: n=8, 11, 8, 10, 10, 9, 10. (N) Change in ovipositor plate length and width (bold and pale colours, respectively) over time, in *D. melanogaster* (blue) and *D. suzukii* (red). Number of samples at the 4 time points in chronological order, in *D. melanogaster*: n=10, 10, 10, 10; in *D. suzukii*: n=10, 10, 9, 10. In all graphs, error bars represent the standard error of the mean.

In summary, the closely matched timing of several, distinct morphological changes, and the dynamics of *senseless* expression, provide multiple lines of evidence that there is no major difference in the rate of ovipositor development between the two species. This means that we can meaningfully compare the same developmental stage in ovipositor development using absolute age (expressed as hours after puparium formation) as an indicator.

With criteria for demarcating and comparing the ovipositor, we can now systematically track the ovipositor size during metamorphosis in both species (Fig. 2M). From 18 to 30 h APF, the presumptive ovipositor remains at approximately the same area in both species. In the initial period of ovipositor elongation, from 30 to 48 h APF, the ovipositor area steadily increases at a comparable rate in both species. Strikingly, however, in the following period from 48 to 54 h APF, while the ovipositor area expands by ~50% in *D. melanogaster*, it expands three times as much by ~150% in *D. suzukii*. Our immunostaining protocol stops working after 54 h APF, presumably due to adult cuticle deposition, preventing the examination of later pupal time points. However, from adult measurements, we infer that, although the ovipositor area continues to increase after 54 h APF in both species, it does so at a very similar rate (by ~20% from 54 h APF to adult) (Fig. 2M). Therefore we conclude that the interspecific difference in ovipositor size is generated in a limited time window, between 48 and 54 hours in pupal development, and is subsequently maintained through later development and into the adult.

Alongside area, we also measured the length and width of the ovipositor when it starts to adopt the form of a blade (Fig. 2N). While ovipositor width modestly increases over development (by ~10% in *D. melanogaster* and ~20% in *D. suzukii*), ovipositor length increases substantially (by ~140% in *D. melanogaster* and ~260% in *D. suzukii*). Hence, within a given species, the increase in ovipositor area during development is chiefly due to an increase in length (rather than width), thus transforming the shape of the tissue with time. Furthermore, between species, the differential increase in ovipositor area is driven predominantly by a differential increase in ovipositor length, thus also explaining the interspecific difference in ovipositor shape. We therefore focused our attention on the cellular parameters that could explain the elongation of the ovipositor.

### Cell proliferation dynamics and total cell numbers are very similar between *D. melanogaster* and *D. suzukii*

We asked whether the ovipositor size difference could be explained by interspecific divergence in cell proliferation or morphology, or both. We first followed cell proliferation dynamics from 6 to 54 h APF. For this, we calculated a mitotic index – the proportion of dividing cells -using an antibody specific to a phosphorylated form of histone H3 (PH3) that labels cells in mitosis (25). In both species, there is a final burst of cell division in the ovipositor from approximately 12 to 36 h APF (Fig. S2A). This is broadly consistent with the timing of terminal waves of cell proliferation described in other imaginal tissues during metamorphosis in *D. melanogaster* (Fristrom and Fristrom 1993). Importantly, there is no major difference in the duration of this burst, nor in the rate of cell division during this period, between the species (Fig. S2A). Furthermore, this shows that cell division in the presumptive ovipositor has stopped at least ~12 hours before the key time window when the interspecific size difference emerges. We can therefore exclude a contribution to the differential ovipositor elongation from processes such as oriented cell division or spatial regulation of mitotic density that have been documented in other systems (12, 26). To investigate further the connection between cell behaviour and ovipositor growth, we needed a method to examine all cells across an entire ovipositor plate, at a given time point. In order to delineate cell contours, we used different markers for adherens junctions in the two species. We used DE-cadherin::GFP (hereafter E-cad::GFP) in *D. melanogaster (27)* and anti-β-catenin antibody staining in *D. suzukii*. These markers allow us to segment almost every cell across an entire ovipositor plate (Fig. S3), facilitating the extraction of several quantitative cell parameters.

With this dataset we first quantified total cell number. Combining this with the cell number estimates we made for stages at 18 to 30 h APF, we find that the pattern of change in ovipositor cell number over time is almost identical in the two species (Fig. 3A), in agreement with the direct measurements of the mitotic index. Moreover, in both species, total cell number actually declines by ~30% from 36 to 54 h APF. In any event, the rate of decline of cell number is comparable in the two species and, ultimately, we find no significant difference in cell number at 54 h APF – the stage at which ovipositor area is ~1.7 fold different between the species (*D. melanogaster* = 1619±45 cells; *D. suzukii* = 1594±51 cells; Student’s t-test p > 0.05). We conclude that differences in total cell number cannot explain the difference in ovipositor length. We therefore turned our attention to cell size.

**Fig. 3.**
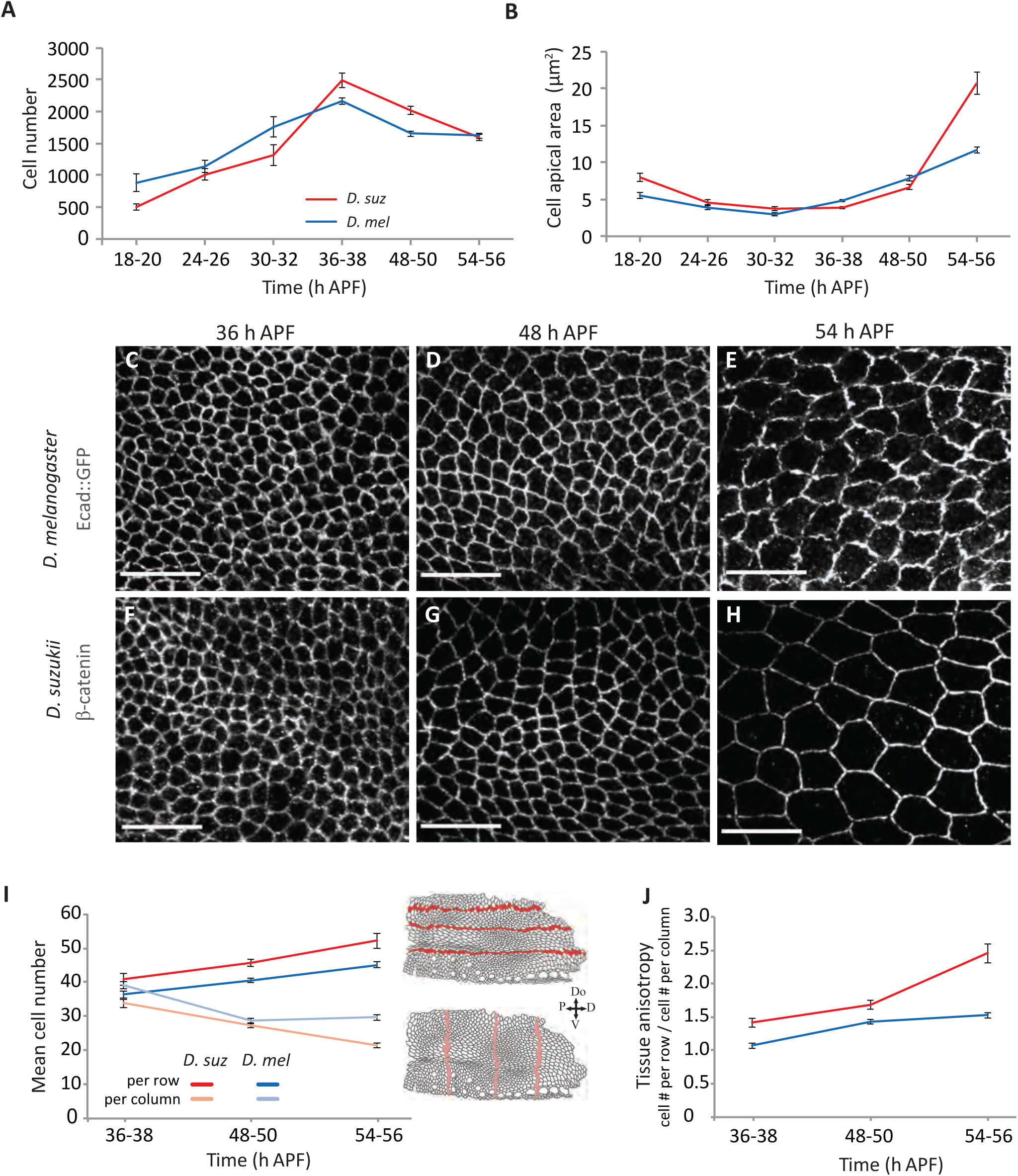
The evolutionary divergence in ovipositor size and shape is driven by accelerated cell size expansion and cell reorganization. (A) Change in mean, total ovipositor plate cell number and (B) in mean cell apical area during pupal development, in *D. melanogaster* (blue) and *D. suzukii* (red). Number of samples at the 6 time points in chronological order, in *D. melanogaster*: n=8, 13, 9, 10, 10, 10; in *D. suzukii*: n=8, 11, 8, 10, 10, 9. (C-H) Illustrative examples of cells in the developing ovipositors of (C-E) *D. melanogaster* and (F-H) *D. suzukii*, at 3 time points during ovipositor elongation. *D. melanogaster* Ecad::GFP, stained for GFP, and *D. suzukii* wild type, stained for β-catenin, to reveal cell apical membranes, shown in grey. All images are to the same scale; scale bar is 10 μm. (I) Change in the mean number of cells per row and per column (bold and pale colours, respectively), in *D. melanogaster* (blue) and *D. suzukii* (red). Drawings on the right show the outlines of segmented cells for an entire ovipositor plate from a particular *D. suzukii* sample at 48 h APF. Selected rows and columns are highlighted in bold and pale red, respectively, illustrating how the average row and column cell number were estimated for a single plate. (J) Changes in tissue shape anisotropy (ratio of cell number per row divided by the number of cell per column) in *D. suzukii* and *D. melanogaster*. For (I, J) number of samples at the 3 time points in chronological order, in *D. melanogaster*: n=10, 10, 10; in *D. suzukii*: n=10, 10, 9. In all graphs, error bars represent the standard error of the mean.

### The evolutionary difference in ovipositor size is driven by accelerated expansion of cell apical area

We next examined if changes in mean cell apical area over time could account for the ovipositor size difference (Fig. 3B). We find that initially, from 18 to 30 h APF, cell size roughly halves in both species. This period coincides with the final wave of cell division in the tissue (Fig. S2A), and it is likely that these divisions are responsible for the decline in cell size – a phenomenon also observed in pupal wing development (28).

However, from 36 h APF, after the cessation of cell division, apical cell surface starts to increase. Indeed we find a striking temporal agreement between the changes in ovipositor area and cell apical area (Fig. S2B, C). From 36 to 48 h APF, we see a comparable rate of expansion in cell apical area in both species, by ~70% over the 12 hours (Fig. 3B; compare panels C, D and F, G in Fig. 3). In stark contrast, from 48 to 54 h APF, while cells continue to expand in *D. melanogaster* by ~50%, cells in *D. suzukii* expand by more than 200%. This represents a dramatic difference in expansion rate of more than four fold. By 54 h APF, an almost two-fold difference in cell apical area has been generated that underpins the interspecific difference in ovipositor size (Fig. 3B; compare panels E and H in Fig. 3). Importantly, we find no significant difference in mean cell size in pupal wings at 54 h APF between the species (Fig. S2D). Hence, the more pronounced apical expansion of cells is not a general feature of *D. suzukii* development, but rather seems to be specific to its ovipositor. It has been suggested that a common mechanism for cell size expansion is polyploidy (29), which can be measured using nuclear size (30). However, in both species, we find no increase in nuclear area from 48 to 54 h APF (Fig. S2E). Hence, we find no evidence that polyploidy can explain the amplified cell size expansion in *D. suzukii*.

Given the substantial increase in cell apical area from 36 to 54 h APF, we wondered whether this expansion in size was uniform, thus preserving cell shape, or whether it was anisotropic, thus stretching the cells. We measured cell shape by fitting an ellipse to each segmented cell and calculating the ratio of the long and short axis of the fitted ellipse. To our surprise, the considerable expansion in cell apical area from 36 to 54 h APF occurs essentially in an isotropic manner in both species. We find only minor and transient changes in apical cell shape during the period of expansion (Fig. S2F). Furthermore, if anything, at 54 h APF the cells are slightly more elongated in *D. melanogaster* than in *D. suzukii*. Therefore although the exaggerated, isotropic cell size expansion in *D. suzukii* accounts for a fraction of the difference in ovipositor size between the species, it cannot account for the change in ovipositor shape during development.

### The evolutionary difference in ovipositor shape is driven by anisotropic reorganization of the tissue

To reconcile the uniform apical expansion of the cells and the elongation of the ovipositor, we examined two further processes: cell orientation and cell reorganization. Given that the ovipositor cells are moderately stretched in both species (mean aspect ratio of ~1.5-1.6 from 36 to 54 h APF; Fig. S2F), we can analyse the orientation of a cell’s long axis with respect to the proximo-distal (PD) axis of the ovipositor. We scored a cell as aligned with the PD axis if the angle created between the cell’s long axis and the ovipositor’s PD axis was less than 45°. In both species, we find a very similar proportion of cells (~50-55%) that have their long axis aligned with the PD axis of the ovipositor at 54 h APF (Fig. S2G). This proportion increases transiently at 48 h APF, specifically in *D. suzukii*, and is concomitant with the transient elongation of cell shape (see above). This suggests that the developing ovipositor is experiencing a stronger deformation around 48 h APF in *D. suzukii* than in *D. melanogaster*. This is only transient, however, and therefore changes in the global pattern of cell orientation cannot explain either the ovipositor elongation during development, or the interspecific differences in shape.

Next we considered the ovipositor anisotropy by examining the organization of the cells along the two major axes of the ovipositor – namely, the PD axis, corresponding to the long axis of the ovipositor (that we call rows), and the orthogonal dorso-ventral (DoV) axis, corresponding to the short axis of the ovipositor (that we call columns) (see diagrams in Fig. 3I). We calculated the average number of cells per row and per column and plotted their dynamics over time (Fig. 3I). The tissue shape anisotropy is then calculated by finding the ratio between the mean number of cells per row and the mean number of cells per column (Fig. 3J).

In both species, the mean number of cells per row increases by approximately 25% from 36 to 54 h APF (Fig. 3I; *D. melanogaster* 36±1 to 45±1 cells; *D. suzukii* 41±2 to 52±2 cells; Student’s t-test p < 0.01). This suggests that, within each species, the increase in cell number along the PD axis contributes to the ovipositor elongation during development. Regarding the between-species difference, from 36 h APF, there is a consistent, small but significant difference in the mean number of cells per row between *D. melanogaster* and *D. suzukii* (at 54 h APF, *D. melanogaster* = 45±1 cells; *D. suzukii* = 52±2 cells; Student’s t-test p < 0.05). Therefore, this ~15% difference in mean cell number per row makes a further contribution to the interspecific difference in ovipositor length.

In contrast to ovipositor length, ovipositor width remains largely constant throughout development in both species (see Fig. 2N). Interestingly, from 36 to 54 h APF, the mean number of cells per column declines in both species -by ~24% in *D. melanogaster* and by ~37% in *D. suzukii* (Fig. 3I). This observation makes sense of the changes in ovipositor shape over time (Fig. 2N). It shows that even though the cell size expansion is isotropic, there is a compensatory reduction in the number of cells along the width of the ovipositor. Overall, the opposing actions of the uniform expansion in cell size and the reduction in column cell number largely cancel out, thus generating only modest changes in width at the same time as substantial changes in length. These changes in the tissue organization increase its anisotropy – and therefore its length – during development both in *D. suzukii* in *D. melanogaster*. However, the anisotropy increase is enhanced in *D. suzukii* compared to *D. melanogaster*, resulting in a longer ovipositor (Fig. 3J).

This reorganization of the cells in the tissue, and in particular the increase in the mean number of cells per row, must result from cell intercalations, at least in part, since there is no more cell proliferation after 36 h APF. Consistent with this idea, in both *D. melanogaster* and *D. suzukii*, we observe a comparable proportion of 4-way vertices (Fig. S2J), indicative that cells shift their relative positions (13). This suggests that cell intercalation increases the number of cells per row, while diminishing the number of cells per column, which results in net elongation of the tissue along the PD axis. Additionally, this overall tissue reorganization is occurring in parallel with a global reduction in cell number. It is therefore possible that this reduction also contributes to the sculpting of the shape of the ovipositor.

### The differences in cell size and tissue shape anisotropy are quantitatively sufficient to explain the evolutionary divergence in ovipositor length

Having characterized several cellular parameters of the developing ovipositors of *D. melanogaster* and *D. suzukii*, we wanted to describe in more precise, quantitative terms, the extent to which these factors can account for the observed, measured ovipositor length differences. We used a simple model for the ovipositor and derived a mathematical equation to calculate the ovipositor length based on the total number of cells, the tissue anisotropy, the cell apical area, and the cell shape (see Materials and Methods in Supplementary Information for further details). Focussing on the 54 h APF time point, we compared the model length estimate with the true, measured value for each species (Fig. 4A). We then introduced coefficients into our mathematical equation to model the extent to which the measured between-species differences in cell parameters are sufficient to transform the *D. melanogaster* into the *D. suzukii* ovipositor, and to assess their respective contribution to length divergence.

**Fig. 4.**
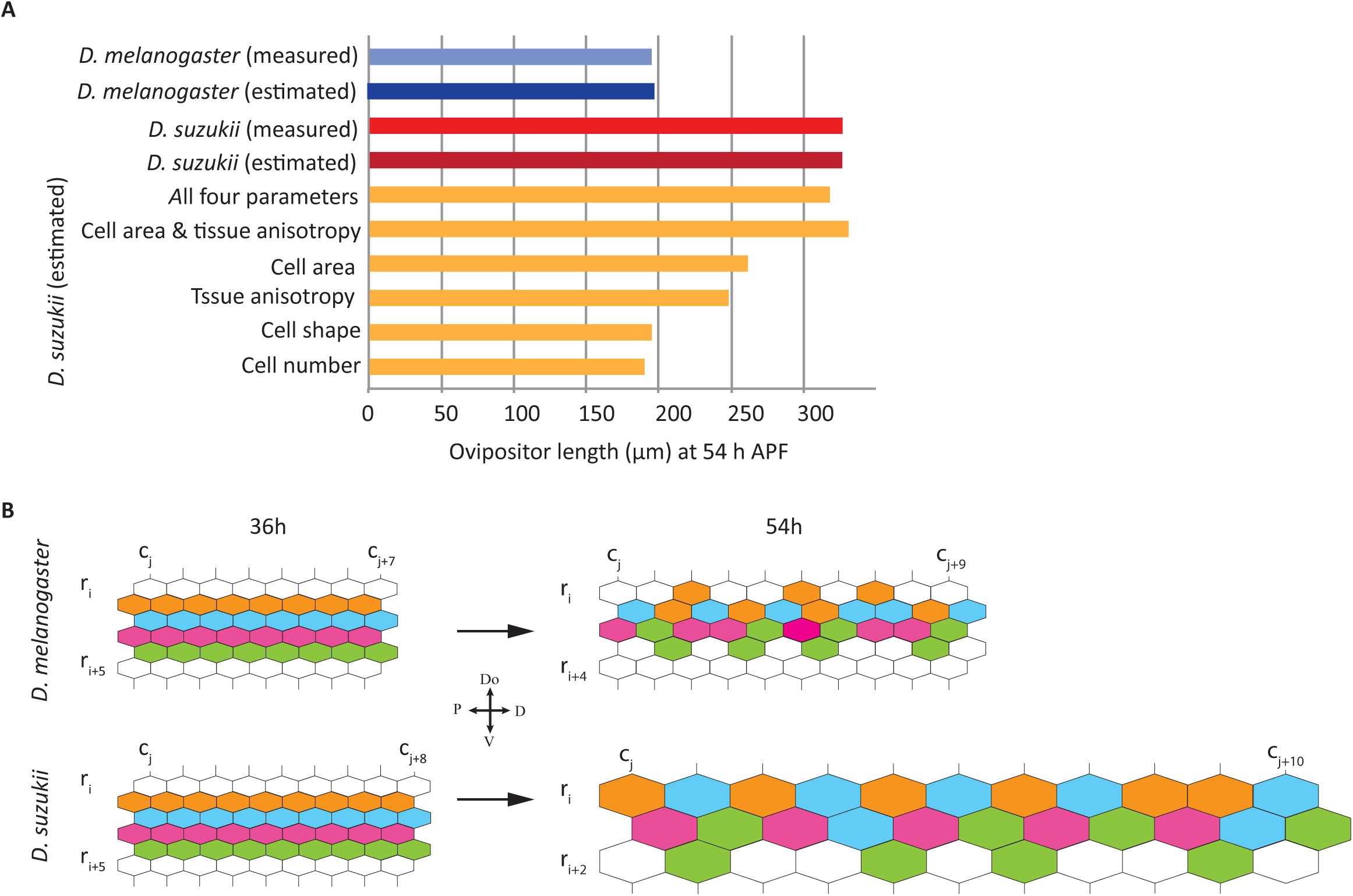
Modelling the development and evolution of ovipositor length shows that the measured differences in cell size and tissue anisotropy are sufficient to explain the bulk of the evolutionary divergence. A. Different model estimates for ovipositor length compared with the true, measured value for ovipositor length in *D. melanogaster* (blue bars) or *D. suzukii* (red bars) at 54 h APF. The *D. melanogaster* (dark blue bar) or *D. suzukii* (dark red bar) estimates are calculated using the species-specific, measured cellular parameters. To assess the relative contribution of the different cellular parameters to the evolutionary divergence, the *D. suzukii* length estimates (yellow bars) are calculated using the *D. melanogaster* values corrected with transformation coefficients for either: i) all four parameters; ii) cell area and tissue anisotropy together; or iii) each cellular parameter in isolation.
B. Schematic representation of the cellular changes that drive ovipositor elongation, in *D. melanogaster* (upper panel) and *D. suzukii* (lower panel), during pupal development from 36 to 54 h APF. Cells are idealized as hexagons and the ovipositor tissue as a hexagonal tessellation, oriented with the proximo-distal (P-D) axis running left-to-right and the dorso-ventral (Do-V) axis running top-to-bottom. Rows and columns are arbitrarily labelled to reflect the change in their number during development, and their difference between species. The schematic illustrates the increase in tissue anisotropy through the increase in the number of cells per row and the concomitant reduction in the number of cells per column during elongation, which acts to balance the isotropic expansion in cell area, thus reducing the net growth in ovipositor width. Colored cells mark neigbhoring cells in a particular row at 36 h APF. By 54 h APF a substantial fraction of the cells have intercalated with one another, contributing further to the tissue elongation.

We find that the estimated ovipositor lengths at 54 h APF (196.8 μm or 326.6 μm) are in very good agreement with the measured values (196 μm or 327 μm), for both *D. melanogaster* and *D. suzukii*, respectively (Fig. 4A). This indicates that our model, although fairly simple, contains sufficient quantitative information to estimate the correct ovipositor length. In addition, applying the coefficient transformation to the *D. melanogaster* parameter values yields an estimated length of 318.2 μm, which closely matches the real value measured in *D. suzukii* (the difference is only 2.6%) (Fig. 4A). This agreement suggests that our mathematical equation accurately captures all the parameters, and their respective quantitative changes, that account for the transformation of the ovipositor length between *D. melanogaster* and *D. suzukii*. Furthermore, our analysis shows that, together, changes in cell apical area and tissue anisotropy are largely sufficient to account for the between-species ovipositor elongation (the difference between the estimated and the measured length is only 1.5%) (Fig. 4A). The differences in cell shape and total cell number, in comparison, only have minor effects on the length divergence between species. In conclusion, the quantitative changes in two cellular parameters, namely cell apical area and cellular rearrangements in the tissue, explain most of the difference in ovipositor length between *D. melanogaster* and *D. suzukii*. If we have missed an additional explanatory factor, it can only be making a marginal contribution to the evolutionary divergence.

## Discussion

D’Arcy Thompson’s goal was to describe mathematically the changes in the size and shape of body parts during development, and between species during evolution. Although his work was descriptively powerful, it was divorced from the underlying cellular processes that sculpt the particular sizes and shapes of morphological structures. With this in mind, our goal in this work was to identify which cellular, morphogenetic processes are responsible for the quantitative changes in a morphological structure between closely related species, and to describe these morphological changes with a mathematical model that reflects the underlying, divergent cellular processes.

We used as a model trait the ovipositor of *D. suzukii*, whose length is almost double that to *D. melanogaster*. We find that most of this divergence in size and shape is generated in a relatively narrow time window, between 48 and 54 hours in pupal development. Furthermore, we find that two shared, cellular processes – namely, the rate of expansion of cell apical area and the reorganization of the cells in the tissue that sets its anisotropy – have quantitatively diverged between the species (Fig. 4B). Finally, we demonstrate with a simple mathematical modelling that the quantitative differences in these cellular processes are sufficient to account for the bulk of the evolutionary divergence in ovipositor length.

First, we find a difference in the rate of cell size expansion. After cell division stops, the ovipositor area continues to grow through isotropic cell apical area expansion. In the key time window, the rate of expansion is accelerated more than four-fold in *D. suzukii*. Ultimately, this produces an almost two-fold difference in cell apical area between the species, which is responsible for the bulk of the difference in ovipositor size. Possible mechanisms for the cell size expansion include cell shape change (for example, cell flattening as observed during dorsal closure in the *Drosophila* embryo (31)), or increase in cell volume. Clarification of this mechanism is a priority for future work. We note that cell surface area also expands in the *D. melanogaster* pupal wing, starting at roughly the same time point (~32 h APF (32). This suggests that cellular expansion during late pupal life might be a developmental feature shared by several body parts.

Second, we find a significant reorganization of cells along the major axes of the ovipositor between the species, resulting in an increased tissue anisotropy. From 36 h APF onwards, there are approximately 15% more cells arranged along the length of the ovipositor in *D. suzukii* than *D. melanogaster*, which makes an additional contribution to the interspecific length divergence. Alongside this, in both species, we documented a significant reduction in cell number along the orthogonal tissue axis (i.e. width) during development. This means that the isotropic cell size expansion is balanced by a reduction in cell number along the ovipositor’s width (see schematic in Fig. 4B). This balancing mechanism allows significant growth to occur along one tissue axis, while producing only modest growth along the orthogonal axis. In *D. melanogaster*, an analogous compensatory mechanism has been described that accounts for anisotropic growth in the wing disc (albeit, it is achieved through a different mix of cellular processes), and thus this may represent a generic morphogenetic strategy for tissue elongation (33).

Several mechanisms might be responsible for the reorganization of the cells in the tissue. The increase in cell number per row, in the absence of cell divisions, indicates that cell intercalations play a role. This reorganization of the cells within the tissue, akin to a convergent extension process, contributes to the increase in ovipositor anisotropy. In addition, within each species, we noticed a striking agreement in the magnitude and timing of the changes in total cell number and in mean cell number per column (Fig. S2H, I). This suggests a second possibility that the global decline in cell number during ovipositor elongation might be driven, at least in part, by spatially patterned apoptosis or cell extrusion that depletes more cell rows than cell columns. In principle, such a mechanism could cause a reduction in the column cell number, though it cannot account for the increase in cell number per row. We note that these hypotheses are not mutually exclusive. One limitation is that we have only been able to examine one face of a given ovipositor plate. Hence we cannot rule out that, as a result of rearrangement or movement, some cells, rather than dying, roll around the contour of the plate and so move beyond our field of view. Clarification of these issues will require creating transgenic fluorescent reporters in *D. suzukii* and establishing live imaging to compare with *D. melanogaster* ovipositor development.

In previous work examining between-species differences in body part morphology at the cellular level, a diversity of cellular processes has been implicated – from changes in cell shape and patterns of cell rearrangement (34), to precise changes in spatial patterns of cell proliferation (12), to joint contributions from both cell number and size (11). A similarly mixed picture emerges from work on the convergent evolution of body size in drosophilids, with the relative contribution of cell number and size varying significantly depending on the specific geographic population or species studied (35 and references therein). In all, it is not yet clear if there is any wider evolutionary significance to the use of different cellular processes in different tissues and species. One hypothesis is that it is the tissue size and shape that are the direct targets of selection, not the cells themselves, and thus the precise cellular processes that are modified will depend on the initial genetic variation in a population, or on mutations that arose, during the period of selection and diversification (36, 37).

Variation in the relative size of morphological traits is a predominant theme of animal morphological evolution. We submit that the fine-tuning of shared cellular processes, similar to the one we have identified in this work, can explain the differential growth underpinning evolutionary changes in body proportion.

## Materials and Methods

They are presented in the Supplementary Information

## Acknowledgments

We thank M. Akam, L. Arnoult, P.F. Lenne and A. Shingleton for comments on the manuscript; N. Walunjkar and A. Lewis for technical help; H. Bellen for the anti-Sens antibody; and B. Aigouy for technical advice and discussions. JEG acknowledges funding support from the EMBO Long-Term Fellowship programme (ALTF 286-2015). The project was supported by the European Research Council under the European Union’s Seventh Framework Programme (FP/2007-2013) / ERC Grant Agreement n° 615789 (BP) and the A^∗^MIDEX project “SuzIDEX” (n° ANR-11-IDEX-0001-02), funded by the “Investissements d’Avenir” French Government programme, managed by the French National Research Agency (ANR) (BP). Imaging was performed on the PiCSL-FBI core facility (IBDM, AMU-Marseille) supported by the French National Research Agency through the “Investissements d’Avenir” programme (France-BioImaging, ANR-10-INBS-04).

